# Selection shapes the landscape of functional variation in wild house mice

**DOI:** 10.1101/2021.05.12.443838

**Authors:** Raman Akinyanju Lawal, Uma P. Arora, Beth L. Dumont

**Affiliations:** The Jackson Laboratory, 600 Main Street, Bar Harbor, Maine 04609, USA; Tufts University, Graduate School of Biomedical Sciences, 136 Harrison Ave, Boston MA 02111

**Keywords:** Genetic diversity, *Mus musculus*, commensalism, genetic disorder, Mendelian disease, adaptation, positive selection, evolution, amylase, metabolism

## Abstract

**Background:** Through human-aided dispersal, house mice have recently colonized new and diverse habitats across the globe, promoting the emergence of new traits that confer adaptive advantages in distinct environments. Despite their status as the premiere mammalian model system, the impact of this demographic and selective history on the global patterning of disease-relevant trait variation in wild mouse populations is poorly understood.

**Results:** Here, we leveraged 154 whole-genome sequences from diverse wild house mouse populations, subspecies, and species to survey the geographic organization of functional variation and systematically identify signals of positive selection. We show that a significant proportion of wild mouse variation is private to single populations, including numerous predicted functional alleles. In addition, we report strong signals of positive selection at numerous genes associated with both complex and Mendelian diseases in humans. Notably, we detect a significant excess of selection signals at disease-associated genes relative to null expectations, pointing to the important role of adaptation in shaping the landscape of functional variation in wild mouse populations. We also uncover strong signals of selection at multiple genes involved in starch digestion, including *Mgam* and *Amy1*. We speculate that the successful emergence of the human-mouse commensalism may have been facilitated, in part, by dietary adaptations at these loci. Finally, our work uncovers multiple cryptic structural variants that manifest as putative signals of positive selection, highlighting an important and under-appreciated source of false-positive signals in genome-wide selection scans.

**Conclusions:** Overall, our findings underscore the role of adaptation in shaping wild mouse genetic variation at human disease-associated genes. Our work highlights the biomedical relevance of wild mouse genetic diversity and unsdercores the potential for targeted sampling of mice from specific populations as a strategy for developing effective new mouse models of both rare and common human diseases.

## Introduction

House mice (*Mus musculus*) are the premier mammalian model system for biomedical research. However, as a consequence of their unique origins from a small pool of founder animals [1], classical inbred mouse strains capture a limited subset of the genetic variation found in the wild mouse populations [2, 3]. Indeed, inbred mice form a monophyletic group within *Mus musculus* [2]. Additionally, at >97% of genomic loci, genetic variation across inbred mice can be reconciled into fewer than ten distinct haplotypes [1]. Thus, inbred mouse genomes harbor numerous “blindspots” over which there is limited genetic diversity that can be linked to phenotypic variation. Further, due to their history of selective breeding for traits of interest and outcrossing between divergent house mouse subspecies, the complex multiallelic nature of trait variation in current panels of inbred strains may not faithfully model complex trait architecture in natural populations, including humans [4].

Wild house mouse genomes represent a largely unexplored reservoir of potential disease-associated genetic variation. Several lines of evidence serve to powerfully illustrate this unrealized potential. First, wild-derived inbred mice, which capture natural variation in a fixed, inbred state, are commonly outliers in strain surveys of disease-related phenotypes [5]. Second, a recent exome sequence analysis of a panel of 26 wild-derived inbred strains identified 18,496 non-synonymous variants that are not segregating among common classical inbred strains [2]. Although the phenotypic effects of these variants are not known, many are undoubtedly functional. Finally, phenotypic surveys of wild-caught house mice have already uncovered significant variation in multiple disease-associated traits, including body mass, metabolism, and behavior [6, 7].

Although wild mice harbor increased genetic variation relative to the classical inbred strains, the population genomic organization and global distribution of wild mouse diversity remain largely unknown. In humans, a significant body of genetics research has underscored the role of adaptation in shaping global patterns of diversity, including variants linked to disease risk and incidence [8]. For example, alleles that conferred a survival advantage to ancient humans during times of starvation have been linked to metabolic disorders in contemporary, food-secure modern human societies [9]. The evolution of malaria resistance has also led to high rates of sickle cell anemia in certain human populations [8, 10]. Similarly, many genes associated with the adaptive evolution of the human brain are linked to neuropsychiatric and neurodevelopmental diseases, including autism and schizophrenia [11-15]. In contrast, the extent to which natural selection may have shaped genetic diversity and disease suceptibility in wild house mice has not been thoroughly explored.

House mice are a species complex composed of three principle subspecies that diverged from a common ancestral population on the Indian subcontinent ∼500,000 years ago [16]. *Mus musculus castaneus* is endemic to Southeast Asia. The native range of *M. m. musculus* extends from Eastern Europe to Northern Asia. *M. m. domesticus* is native to the Middle East and Western Europe. Approximately 10,000 years ago, *M. musculus* developed a commensalism with human agricultural societies. This ecological transition was likely accompanied by dietary shifts, changes in environmental pathogens, and the emergence of new behaviors. Through human-aided dispersal over the last ∼10,000 years, *M. musculus* have expanded their home range to Africa, Australia, and the Americas. This incredible and recent geographic expansion required further local adaptation to multiple distinct ecosystems, including arid, high-altitude, cold, and extreme heat environments, as well as exposure to new pathogens. Adaptation to these new environmental pressures has potentially left unique and detectable footprints in patterns of genomic diversity across contemporary wild mouse populations.

To evaluate the impact of local adaptation and population history on the global patterning of putatively functional wild mouse genetic variation, we analyze a set of 154 publicly available diverse wild house mouse genome sequences in an evolutionary framework. We profile the global organization of predicted functional variants across multiple populations from each of the three core house mouse subspecies and perform genome-wide scans for positive selection to assess the role of adaptation in shaping patterns of genetic diversity across these populations. Overall, our study reveals the landscape of functional variation in wild house mouse populations and underscores the promise of targeted sampling of mice from specific populations and environments as a strategy for developing new models of both rare and common human diseases.

## Results

### Wild house mice capture significant, and potentially functional, diversity that is absent from inbred laboratory mice

We utilized 154 publicly available wild mouse whole-genome sequences for this study [6, 17, 18]. This panel features genome sequences from *M. spretus* and multiple populations from each of the three principle *M. musculus* subspecies: *M. m. domesticus* (4 populations: Eastern United States (America), France, Germany, Iran), *M. m. castaneus* (2 populations: India, Taiwan), and *M. m. musculus* (3 populations: Afghanistan (Afghan), Kazakhstan (Kazakhstani), Czech Republic (Czech)). The combined *Mus* dataset yields ∼154 million biallelic autosomal single nucleotide polymorphisms (SNPs), including 617,156 missense, 7,615 nonsense, and 985,873 synonymous SNPs. Of these, 15,104 SNPs in 6788 unique genes are predicted to be highly deleterious and disrupt gene function. Within *M. musculus* (n = 146 genomes), there are ∼121 million autosomal SNPs, including 772,614 synonymous, 493,090 missense, 6,216 nonsense, and 12,396 highly deleterious SNPs. Consistent with prior work [19], we observed the highest genome-wide nucleotide diversity, π in *M. m. castaneus* (0.0249), followed by *M. m domesticus* (0.0172), and *M. m. musculus* (0.0160). Variant statistics for each population and subspecies are provided in Figures 1a and 1b.

**Figure 1.**
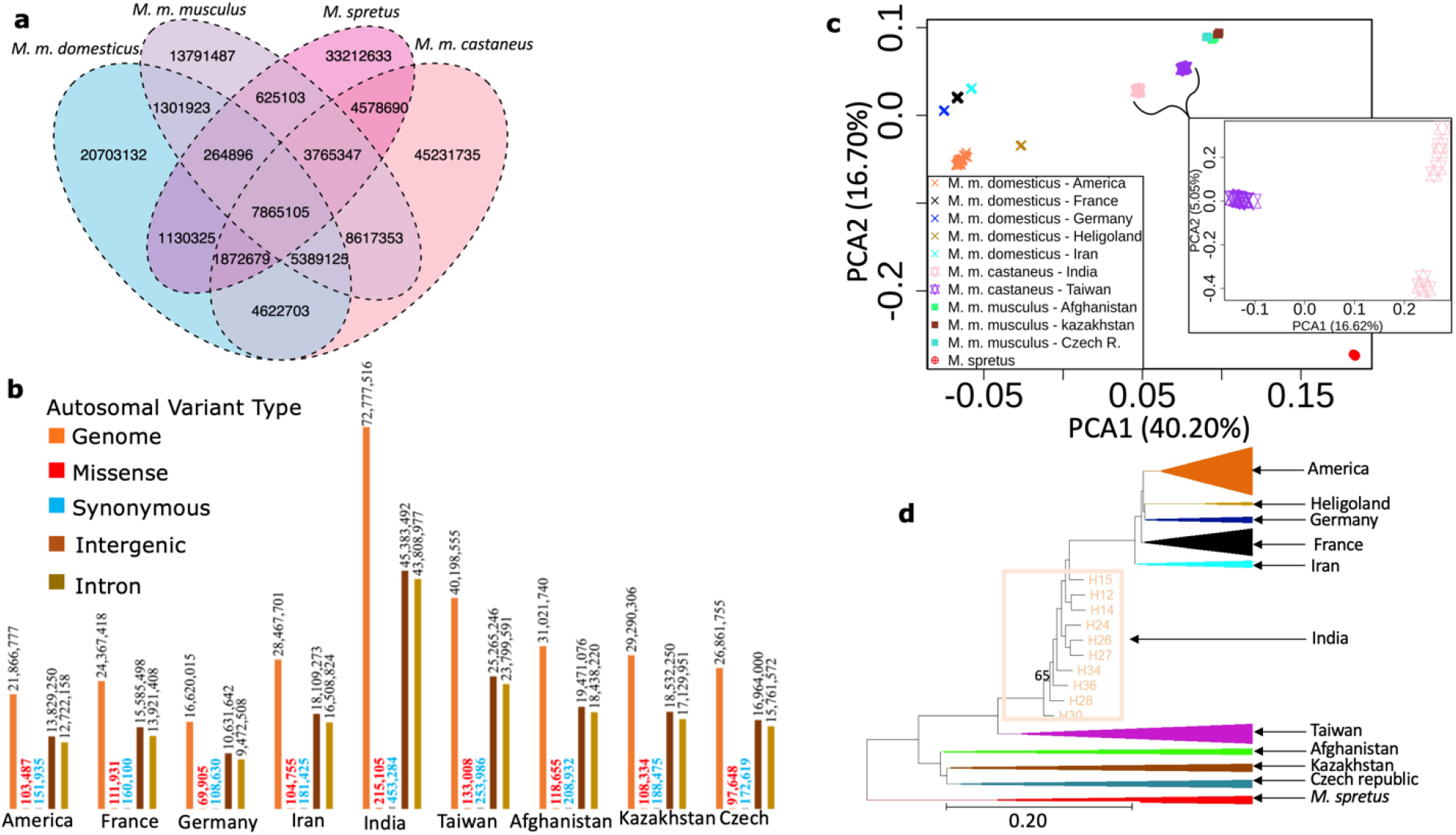
Functional annotation of wild mouse genetic diversity. (**a**) Venn diagram of shared and private SNPs in each house mouse subspecies and species. (**b**) Total numbers of autosomal, missense, synonymous, intergenic, and intronic SNPs in each *M. musculus* population. (**c**) Principal component analysis for all 154 wild mouse genomes. The inset zooms into the two *M. m. castaneus* populations and reveals greater diversity among samples from the Indian population. (**d**) Maximum likelihood phylogenetic tree from all 154 wild mouse genomes. For ease of visualization, samples from most populations are collapsed, with triangle width scaled by the number of samples in that population. One node with <100% bootstrap support is labeled. All other nodes are supported by 100% of bootstrap replicates. The color legend in (**c**) is also applicable to (**d**).

On the X chromosome, we identify an additional 22,716 synonymous, 17,957 missense variants, 367 nonsense, and 654 deleterious mutations across all samples. The vast majorty of these variants are segregating within *M. musculus* populations, and are not private to the outgroup *M. spretus*. Specifically, we find 4,157 (6,537; 4,587) missense, 101 (93, 74) nonsense, 4,216 (10,185; 5,487) synonymous, and 215 (187; 147) deleterious mutations on X chromosome segregating in *M. m. domesticus* (*M. m. castaneus*; *M. m. musculus*), respectively. *M. spretus* has 6,087 missense, 101 nonsense, 8,586 synonymous, and 164 deleterious variants on the X chromosome.

Approximately 69.3% of the autosomal variants in *M. m. domesticus*, 63.9% of *M. m. castaneus* autosomal variants, and 53.7% of *M. m. musculus* autosomal variants are not segregating in panels of common inbred mouse strains. Within *M. musculus*, 13,023 of these wild variants are predicted to be highly deleterious. Although a subset of these variants may be false positives, it is nonetheless evident that wild house mouse genomes harbor substantial unexplored, and potentially functional, variation.

### Patterns of genetic relatedness among wild mouse samples

As our dataset was compiled from multiple prior studies [6, 17, 18], we next examined kinship and relatedness metrics among samples from each population to identify any close relatives. Fourteen pairs of animals have kinship coefficients >0.08, indicating first- or second-degree relatedness. Importantly, we obtain qualitatively identical findings regardless of whether closely related individuals are included or excluded from our analyses (Additional file 1: Table S1). Given the small sample sizes for several of the wild mouse populations and the robustness of our findings to the relatedness among samples, we opt to include all samples in the analyses presented below.

We then performed phylogenetic and principal component analyses (PCA) to assess genetic relationships among populations. As expected, populations from the same subspecies group together in both PCA and phylogenetic analyses (Figures 1c and 1d). We utilized two independently sampled populations from around the Massif Central region in France. There is no clear evidence for genetic stratification of these samples (Figure 1c and 1d), and we combine these two independently sampled populations in our analyses. We observe greater differentiation between *M. m. castaneus* populations from India and Taiwan than between populations within other subspecies. This result is expected given the presumed ancestral origins of house mice on the Indian subcontinent and the large effective population size of this population [20], in contrast to the recent colonization of Taiwan (Figure 1c inset). Further, consistent with these differences in population history [21], genome-wide heterozygosity is markedly reduced in the Taiwanese population [18] compared to the Indian population (4% vs 25%). The American and the Heligoland populations of *M. m. domesticus* are clearly differentiated from those of mainland Europe and Iran (Figures 1c and 1d), underscoring the genetic impact of founder effects during the recent colonization of these regions.

### Predicted functional properties of population-private variants

As a by-product of their unique demographic origins and history of local adaptation from new or low frequency mutations, individual house mouse populations are expected to harbor uniqu suites of private variants, including alleles with effects on fitness. To understand the prevalence and functional impact of such alleles, we identified variants private to each population, limiting our focus to those with a minimum allele count of 2 in the focal population to alleviate the influence of sequencing and genotyping errors. Because of the small sample sizes for each population, we acknowledge the likelihood that many of the variants marked as “private” are potentially present at low frequency in other populations.

Overall, we identify ∼31.7 million population-private autosomal variants, representing approximately 20.6% of all segregating autosomal variants in *M. musculus*. Thus, there is considerable geographic structuring of global mouse genomic diversity. Despite the prominent role of human-facilitated migration and colonization in recent house mouse history [22], individual populations continue to harbor large loads of private variants. As expected based on estimates of effective population sizes and recent demographic histories [20], we find the highest numbers of population private variants in the *M. m. castaneus* populations and in the Iranian *M. m. domesticus* population.

Although most population private variants are in intergenic regions and are likely neutral, an appreciable fraction reside in coding regions where they may exert effects on individual fitness (Table 1). Specifically, we identify 1,483 predicted loss-of-function (LOF) variants in 1,205 unique genes across the nine surveyed *M. musculus* populations. Of special note, we find a private stop-gain mutation at codon position 72 of *Mdm4* (chr1:133,011,141) that is at ∼42% frequency in the Afghanistan population. *Mdm4* is a negative regulator of *p53* and is upregulated in a number of human cancers [23, 24]. Similarly, a private mutation to the German population in *Mutyh* (chr4:116815563; ∼14% frequency) disrupts a splice acceptor site and is predicted to abolish gene function. *Mutyh* is involved in oxidative DNA damage repair and mutations in this gene are associated with hereditary forms of colorectal cancer [25] and biases in the spectra of both germline [26, 27] and somatic mutations [28]. There are currently four knockout and/or targeted mutation mouse models available from commercial vendors for each *Mdm4* and *Mutyh* [29]. Our analyses reveal that organic evolutionary processes have already generated natural loss-of-function alleles for these, and presumably many other, critical disease-related genes.

**Table 1.** Number of coding and predicted functional variants per population.

| Population | Number of Private Variants | Number of synonymous private variants | Number of missense private variants | Number of stop private variants | Number of predicted deleterious variants | Number of predicted LOF variants |
| --- | --- | --- | --- | --- | --- | --- |
| America | 1,025,466 | 8,675 | 9,637 | 155 | 242 | 151 |
| France | 2,208,483 | 10,190 | 11,521 | 221 | 350 | 218 |
| Germany | 545,881 | 2,178 | 2,506 | 51 | 74 | 39 |
| Iran | 3,333,440 | 15,430 | 11,548 | 149 | 235 | 145 |
| India | 16,025,598 | 70,201 | 35,493 | 447 | 745 | 359 |
| Taiwan | 3,917,054 | 19,969 | 15,957 | 244 | 410 | 225 |
| Afghanistan | 1,472,430 | 7657 | 6566 | 124 | 170 | 102 |
| Czech Republic | 1,315,866 | 6582 | 6489 | 112 | 169 | 106 |
| Kazhakstan | 1,872,782 | 9,403 | 8,902 | 174 | 230 | 138 |
| Total | 31,717,000 | 150,285 | 108,619 | 1,677 | 2,625 | 1,483 |

### Detecting signals of positive selection in wild mouse genomes

Just as observed in human populations [8], local adaptation has almost certainly molded the geographic distribution of disease-associated trait variation in wild mice. To directly investigate this possibility, we carried out genome-wide scans for positive selection in each of the nine surveyed wild mouse populations.

Strong positive selection on an adaptive allele will result in its rapid sweep to high frequency or fixation in a population. This process will yield a localized reduction in genetic diversity at the selected site, a signature referred to as a “selective sweep”. The strength of this trademark signal is governed by a complex interplay of population genetic variables, including the magnitude of selection, the initial frequency of the selected allele, and the local rate of recombination.

We employed three population genetic diversity summary statistics – *H*_p_ (pool heterozygosity) [30], π (nucleotide diversity) [31], and Tajima’s D [32] – to identify signals of selection in each of these populations. We computed each statistic in 20 kb sliding windows across the genome using a 10 kb step size. This window size is less than the expected scale of linkage disequilibrium decay in previously surveyed wild mouse populations [3].

A key challenge for the interpretation of genome-wide scans for selection is to distinguish regions truly evolving via positive selection from outliers of the neutral diversity distribution. For example, certain demographic scenarios can induce genome-wide reductions in diversity that may masquerade as pervasive positive selection [33]. One widely used approach to circumvent this challenge is to apply coalescent simulations that realistically model the ancestry of the analyzed sample in order to derive an empirical distribution of the test statistic under the assumption of neutrality. However, the complex history of human-aided migration and introgression among house mice pose significant challenges to the inference of accurate demographic models for each population [20]. For this reason, we instead focus on a hard empirical threshold (the lower 0.1% tail) to ascertain regions evolving via strong positive selection. The precise percentage of the genome that is subject to positive selection is likely greater than this cutoff [34]. However, this conservative, heuristic approach eliminates the need to devise population-specific demographic history models that invoke a large number of assumptions, many of which may not be correct or are potentially misleading. We then collapsed adjacent windows passing this cut-off into contiguous loci to define candidate regions. We focus on regions detected as outliers by the *H*p statistic and by at least one of the other two statistics. Additional files 2 – 4: Figures S1 – S3 display the genome-wide distributions of these three summary statistics in each population.

Overall, we identified 154 unique putative sweep regions across the four *M. m. domesticus* populations, including 30 in the French population, 14 in the Iranian population, 29 in the German population, and 86 in the American population. A total of 172 selective sweep loci were identified in *M. m. castaneus*, with 57 identified in the Indian population and 115 in the Taiwanese population. We uncovered 188 putative selective sweep loci in *M. m. musculus*. Of these, 68 were observed in the population from Afghanistan, 60 in the Kazakhstani population, and 64 in the Czech population. We also identified 63 candidate selective sweeps in *M. spretus*. Additional files 5 – 8: Tables S2 – S5 present comprehensive catalogs of these candidate regions, including shared signals of positive selection between populations.

Positive selection is expected to operate exclusively on functional genomic regions, but there is no *a priori* expectation that neutrally evolving loci should be enriched for functional annotations [35]. Approximately 86% of the selective sweep loci identified in our analysis span at least one protein-coding gene. In contrast, in 1000 independent simulations of random size-matched genomic intervals, at most 37% overlapped protein-coding genes *(p*<0.001). The marked enrichment for protein-coding annotations in our selective sweep windows suggests that our candidate regions are strongly enriched for bonafide targets of positive selection.

### Cryptic structural variation manifests as false-positive signals of selection

We noted that many candidate selective sweep regions overlapped annotated segmental duplications and polymorphic structural variants previously described in laboratory mouse strains. For instance, we observed a sharp decrease in *H*p, π, and Tajima’s D at chr4:112.23 -112.61 Mb, a locus spanning a cluster of paralogs in the *Skint* gene family (Additional file 9: Figure S4a). Relative to the C57BL/6J mouse reference genome, at least 13 inbred strains carry a deletion in this region that spans three paralogs (*Skint3, Skint4*, and *Skint9)* [36, 37]. We analyzed patterns of read depth at the *Skint* locus in our wild mouse samples (Additional file 9: Figure S4b and c) and confirmed that a single deletion allele segregates at frequencies 57%, 80%, and 82% in wild *M. m. domesticus, M. m. castaneus*, and *M. m. musculus* populations, respectively.

These findings raise the possibility that cryptic deletions or other structural variants may commonly lead to local reductions in the number of surveyed haplotypes and an expected, concomitant loss of diversity. Critically, prior studies demonstrate that wild house mouse populations harbor high loads of structural variation [17, 38] which, if ignored, could yield abundant false-positive signals of positive selection. We applied a post-hoc read depth filter to mask regions of the genome present in a non-diploid state (see Methods). After applying this key quality control step, the number of putative selection windows decreased from 607 to 577. Thus, approximately 5% of all regions originally identified in our analysis are likely false-positive signals attributable to structural variation. Our findings underscore the significant impact of cryptic structural variation on genome-wide inference of positive selection, and urge for the implementation of basic filters to eliminate copy number variable regions in QC processing for genome-wide scans. All analyses presented below focus on this refined set of candidate positive selection regions.

### Targets of positive selection in wild house mouse populations

Our catalogs of positive selection emphasize several known and recurrent targets of adaptive evolution in mammals. For example, in the French population of *M. m. domesticus*, we identified a strong selective sweep at *Epas1* (chr17:86.78 - 86.80 Mb). *Epas1* is a transcription factor that is activated under hypoxic conditions and prior studies have linked variation at this gene to high-altitude adaptation in mammals and birds [39, 40]. Mice from this population were collected from the mountainous Massif Central region of France [17], where oxygen levels may be reduced to as much as 81% compared to sea level. We also observed signals of positive selection around *Mgam* (chr6:40.66 - 40.79 Mb) in the Iranian and Afghan populations (Figure 2, Additional files 5 - 6: Table S2 and S3). *Mgam* encodes a starch digestion enzyme and prior work has implicated this gene in the adaptation to starch-rich diets during dog domestication [41] and the transition to agriculture in ancient Andean humans [42]. Similarly, in the French population, we observed positive selection around *Amy1* (chr3:113.52 - 113.56 Mb), an enzyme involved in carbohydrate digestion, confirming findings from a prior selection scan in wild mice [43]. In the Kazakhstani population of *M. m. musculus*, a selective sweep spans *Herc2* and *Oca2* at chr7:56.23 – 56.25 Mb. Genetic variation in *Herc2* is linked to the pigmentation in the skin, hair, and eye, and *Oca2* plays a role in melanin synthesis and eye color determination [44, 45]. Analyes of selection in diverse humans populations have revealed strong selection pressures at this locus [45].

**Figure 2.**
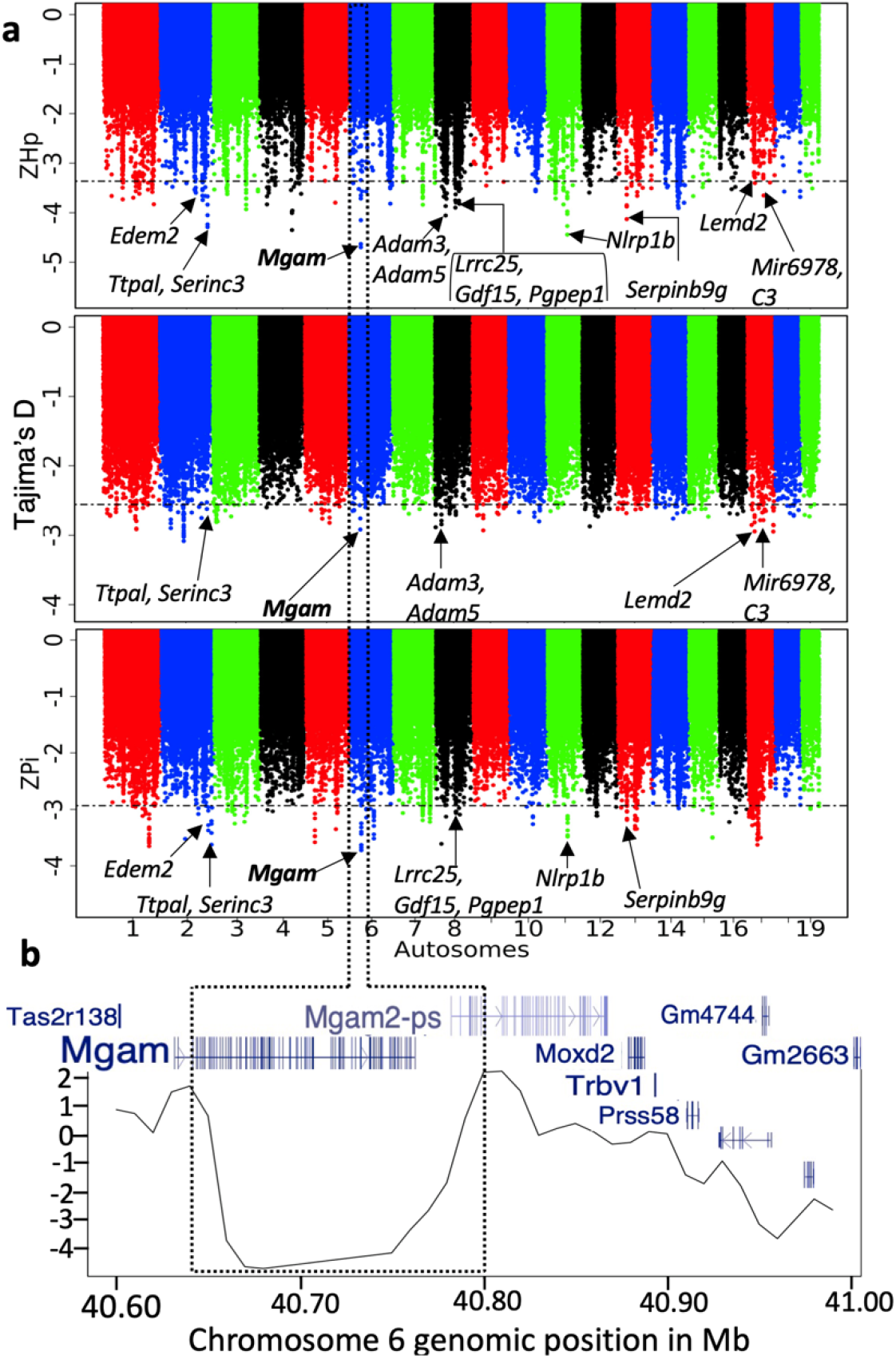
Signatures of positive selection in the Iranian population of *M. m. domesticus*. **(a)** The genomic distribution of *H*_p_,, and Tajima’s D. Horizontal lines on the first three panels correspond to the 0.1% genome-wide empirical significance threshold. Several genes within significant peaks are annotated (also see Additional file 5: Table S2). Panel (**b**) provides a close-up of *H*_p_ across the *Mgam* locus on chromosome 6: 40.66 – 40.79 Mb. The gene track overlaid on panel (**b**) shows that the signal of selection is localized to the coding region of *Mgam*.

In parallel, our genome-wide scans uncovered several regions with no previously reported history of adaptive evolution in mammals. For instance, in the German population of *M. m. domesticus*, we observed a notable peak spanning *Cdan1* and *Ttbk2* on chromosome 2 (120.73 - 120.77 Mb). *Cdan1* functions in chromatin assembly with mutations in the gene linked to congenital dyserythropoietic anemia [46]. *Ttbk2* plays a key role in ciliogenesis, the development of the cerebellum, and balance coordination [47]. We also found evidence for a selective sweep around *Cdan1* in the American population (Additional file 5: Table S2). To our knowledge, our report represents the first evidence for adaptive evolution at this locus, although the specifi environmental pressures that have led to these sweep signals remain to be determined.

In the Indian population of *M. m. castaneus*, we report a strong selective sweep at chr5:104.50 - 104.54 Mb. This locus harbors two unannotated genes, as well as the upstream regulatory region and first two exons of *Pkd2. Pkd2* is expressed in the embryonic and adult kidney and is critical for maintaining the highly differentiated state of the polarized epithelia forming adult renal tubules and bile ducts [48]. The strongest signal of positive selection in the Taiwanese *M. m. castaneus* population localizes to chromosome 1 (159.51 - 159.54 Mb) and spans the 5’ end of a single gene, *Tnr. Tnr* is highly expressed in the central nervous system and prior studies have established its essential functions in cognition and motor coordination [49] (Additional file 7: Table S4).

In each of the three *M. m. musculus* populations, we observe a pronounced selective sweep signal at chr19:3.65 – 3.74 Mb (Additional file 6: Table S3). This locus spans *Lrp5*, as well as several transcripts of unknown function. *Lrp5* has diverse roles in the maintenance of bone mass, eye development, and cholesterol homeostasis [50], and has been implicated in osteoporosis [51]. In the Kazakhstani population, the strongest signal of positive selection resides on chromosome 15 (3.25 – 3.31 Mb) and spans *Ccdc152* and *Selenop. Ccdc152* is poorly studied. *Selenop* encodes a seleno-protein that transports selenium to the plasma, where it is functionally important in thyroid metabolism and protection against oxidative stress [52]. In the Czech population, we identified a strong selective sweep at *Grik3*, an excitability neurotransmitter receptor on chromosome 4 (125.55 - 125.57 Mb).

The most prominent selective sweep signal in *M. spretus* (Additional file 8: Table S5) localizes to chromosome 16 (29.58 - 29.65 Mb) and encompasses a single gene, *Opa1. Opa1* is a dynamin-like GTPase gene that functions at the inner mitochondrial membrane and plays a critical role in visual perception [53]. A second significant peak on chromosome 14 (27.48 - 27.51 Mb) spans *Ccdc66*, a gene implicated in retinal morphogenesis [54].

### Signals of positive selection on the X chromosome

Mammalian sex chromosomes are crucibles of sex-biased selection signals [55]. Our strategy for identifying regions under positive selection does not allow us to define the relative frequency of selection on the X and autosomal genome compartments, but we nonetheless extended our analysis to identify regions evolving via positive selection on the X. Overall, we identified 39 unique X-linked loci putatively under positive selection across the nine populations of *M. musculus* and *M. spretus* (Additional file 10: Table S6). Six of the 39 loci exhibit signals of positive selection in more than one population. For example, a locus spanning *Wnk3*, a serine/threonine protein kinase involved in the regulation of cellular electrolyte homeostasis, has a deficit of diversity in both the American and German populations of *M. m. domesticus*. Populations representing each of the three principle house mouse subspecies – the American, Taiwanese, and Kazakhstani populations – bear similar signatures of positive selection around *Il1rapl2*, a gene involved in central nervous system development and linked to cognitive impairment and disability [56] (Additional file 11: Table S7).

The majority of candidate X-linked positive selection regions overlap a single gene (35/39 = 87.5%). Focusing on the 35 regions with a single presumed gene target of adaptive evolution, we find that most genes (23/35) are expressed in the pre-meiotic cells of testis [57]. In contrast, only two loci harbor genes (*Samt3* and *Sh3kbp1*) that are exclusively expressed in postmeiotic spermatids (Additional file 10: Table S6). The clear enrichment of selection signals at genes expressed prior to the first meiotic division suggests that natural selection has actively sculpted the gene content of the mouse X chromosome, consistent with theoretical predictions and prior work on house mice [58, 59].

### Selective sweeps are enriched for genes implicated in Mendelian diseases and GWAS hits

We noted that many of regions of positive selection in wild mouse genomes overlapped known disease-associated and disease-causal genes in humans (Additional file 11: Table S7). Overall, 43% of sweep windows overlap a gene in the Online Mendelian Inheritan in Man (OMIM) database, a significant increase over random expection (1000 random simulations of size-matched genomic regions: 16% - 38% overlap with OMIM genes). Similarly, 44% of sweep windows overlap a gene in the genome-wide association study (GWAS) catalog (https://www.ebi.ac.uk/gwas/), again in excess of expectations from random simulations (range: 15% - 39% in 1000 simulation replicates).

To investigate these trends on a per-population basis, we estimated the fraction of sweep genes identified in each population that overlap OMIM genes. This quantity varies considerably across the surveyed populations, ranging from 23% - 58% compared to the null expectation from simulation ranging from 21% - 34% (Table 2). *M. spretus* exhibits the highest degree of overlap between the identified putative selective sweep genes and genes in the OMIM database, with the French M. *m. domesticus* population exhibiting the lowest overlap. Similarly, populations vary in the proportion of sweep genes that overlap GWAS hits (25-68%). Taken together, these results suggest that targets of positive selection in the majority of wild mouse populations are significantly enriched for disease-associated genes.

**Table 2:**
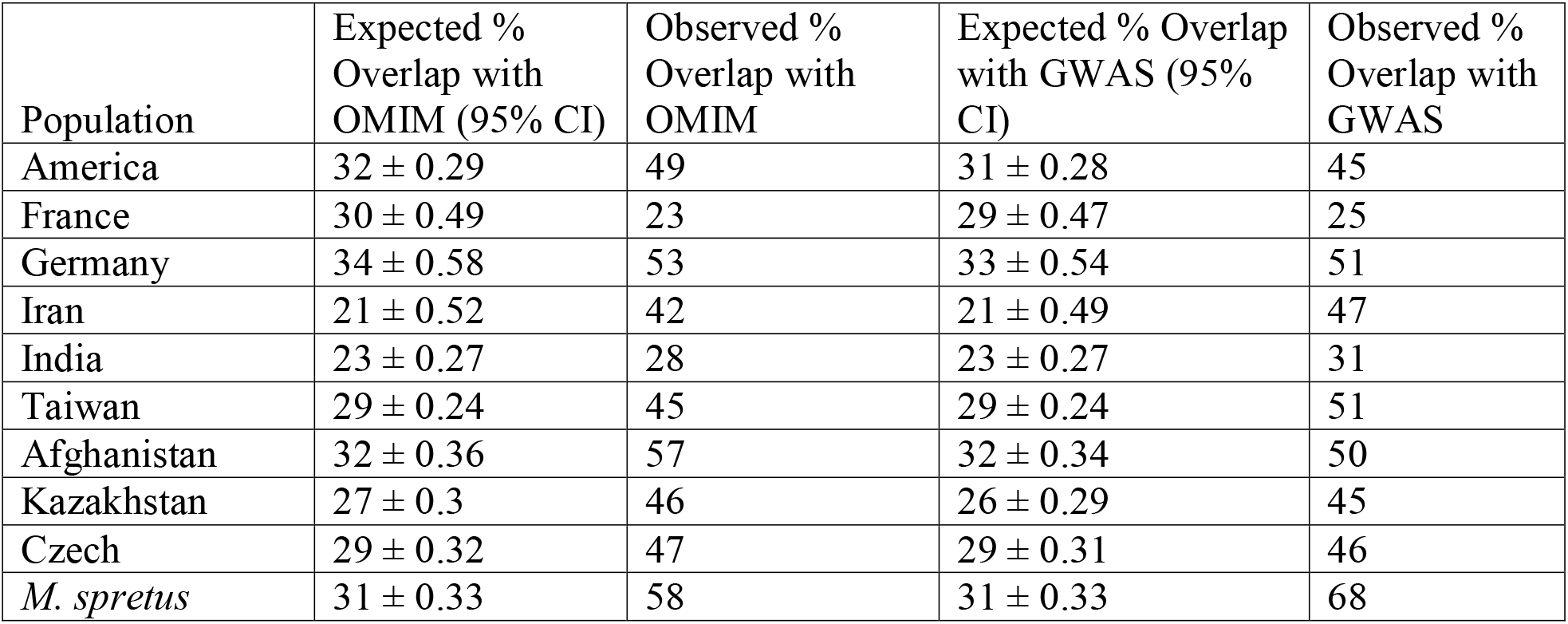
Frequency (%) of genes in selective sweep windows overlapping with genes in OMIM database and GWAS catalog.

### Functional classification of genes evolving via positive selection

We applied Gene Ontology (GO) and Kyoto Encyclopedia of Genes and Genomes (KEGG) enrichment analyses to the sets of selective sweep genes from each population and for all populations within a subspecies. Genes that regulate the immune response to external pathogens are common targets of positive selection across species [60], and we recapitulate this well-known finding. In *M. m. domesticus*, selection targets are enriched for “response to virus” (*Irak3, Hyal2 Fv1, Uri1*, and *Il15*) and “inflammatory response” (*C3, Hyal2, Tusc2, P2rx7, Sema7a*, and *Sgms1*) GO terms. In *M. m. castaneus*, an excess of positive selection genes are annotated to the GO terms “innate immune response” (*Serinc3, Cr2, Herc6, Cr1l*, and *Iglc3*) and “T-cell mediated immunity” (*Cd46*, and *Myo1g*). Similarly, in *M. m. musculus*, there is an enrichment of genes under positive selection annotated to the “immunoglobulin subtype 2” GO term (*Prtg, Mdga2, Nfasc, Pdgfrb, Sema3c, Lrfn5*).

Within *M. m. domesticus*, genes under positive selection are enriched for three KEGG pathways associated with digestion, including “carbohydrate digestion and absorption” (*Mgam, Amy1, Amy2a5, Atp1a4, Atp1a2*), “protein digestion and absorption” (*Col14a1, Atp1a4, Atp1a2*), and bile secretion (*Slco1a4, Atp1a4, Atp1a2*). Enriched functional annotations in *M. m. castaneus* include “Cytochrome P450” (*Ptgis, Cyp2j8, Cyp2c38, Cyp19a*1), “adult feeding behavior” (*A (agouti), Ghrl*), and the KEGG “oxytocin signaling pathway” (*Ppp1r12a, Ryr2, Prkaa1*, and *Mapk7*). Within *M. m. musculus*, genes under positive selection are disproportionately involved in pigmentation (e.g *Vangl1, Oca2, Herc2*), DNA damage (e.g *Mre11a, Casp9, Herc2, Smg1, Hmga2, Tesmin*), and cholesterol metabolism (e.g *Osbpl7, Hsd17b4, Cpt1a, Nr1h5*). Additional files 5 – 8: Tables S2-S5 provide comprehensive summaries of findings from these functional enrichment analyses.

### An initial test of regulatory versus coding mechanisms of adaptive evolution in wild mice

An enduring question in evolutionary biology concerns the relative roles of adaptation on coding versus regulatory sites [61]. Although it is beyond the scope of this article to resolve this long-standing debate, we leveraged published RNA-seq data [17] from a subset of the wild *M. m. domesticus* mice used in these genome-wide selection scans to test for significant differences in gene expression at a handful of selection targets (Additional file 5: Table S2 and Additional file 12: Figure S5). Of note, *Epas1* is under positive selection in the French population and is significantly upregulated in liver and muscle tissues of mice from France as compared to mice from the German population. However, we do not observe differential expression of this gene in the heart, as previously shown for high-altitude adapted deer mice [62]. We find no significant differences in *Mgam* expression levels in digestive tissues (gut, liver) among the *M. m. domesticus* populations, suggesting that positive selection at this locus may act on coding sites that alter enzymatic activity. This finding aligns with the observed *H*_p_ profile across this locus, which reveals a pronounced drop in diversity that is restricted to coding portions of the gene (Figure 2b). *Cry1*, a highly conserved gene implicated in the maintenance of circadian rhythm, shows upregulation across multiple tissues in French mice compared to mice from Germany and Iran, consistent with the signal of adaptation at this locus which is restricted to the French population. Finally, *Amy1* is under positive selection in the French population and is upregulated in gut tissues from both the Iranian and French populations relative to mice from the German population. This finding is consistent with possible regulatory modes of adapative evolution at this locus. Despite the limited scope of this analysis, our findings demonstrate that positive selection has acted on both coding and regulatory sites in wild mouse populations.

## Discussion

Here, we analyzed the genomes of 154 wild-caught mice to assess the population-wide distribution of functional genetic diversity and establish the contribution of positive selection to the global patterning of disease-relevant trait variation. We show that a large fraction of wild mouse variation is specific to individual populations, including numerous predicted loss-of-function variants that could be useful in the context of disease modeling. Further, our work has sythesized a comprehensive catalog of candidate genes and genomic regions evolving via positive selection in diverse wild house mouse populations. Our surveyed populations inhabit distinct environments that differ in altitude, average temperature, aridity, and human population density. These environmental differences have created unique opportunities for population- and subspecies-specific adaptations, including the emergence of adaptive traits that may confer differences in disease susceptibility. Several exciting themes emerge from this catalog.

First, like many other animal species [60], genes involved in immunity and sensory perception are common targets of adaptive evolution in wild house mice. Across populations and subspecies, we identify multiple sweep regions spanning genes with immune-related functions. The diverse suite of pathogens endemic to each population’s environment has likely imposed strong selective pressures on the immune system. We also document positive selection signals at multiple olfactory receptors (ORs). The OR repertoire is known to evolve rapidly, with notable gains and losses across the mammalian tree [63]. Interestingly, we find few shared signals of selection at ORs across wild mouse populations (Additional file 5 – 7: Tables S2 – S4), suggesting that positive selection has led to precise, population-specific OR portfolios tuned to the detection of specific aromatic compounds in the prevailing environment.

Second, several genes that are evolving via positive selection in house mice are also targets of adaptive evolution in human populations. For example, *Epas1* has been implicated in high altitude adaptation in several human populations and we observed a genetic signature of recent selective sweeps at this locus in mice from a mountainous region in France. Similarly, *Mgam* is evolving under adaptive evolution in both an Andean human population [42] and in wild mouse populations from Iran and Afghanistan. These instances of parallel evolution suggest that wild mice could serve as powerful models for dissecting the molecular basis of some adaptative traits in humans.

Third, our study uncovers loci that may have contributed to the development of a successful commensalism between house mice and humans. Recent archeological evidence shows that mice emerged as commensals with humans approximately 14,500 cal. BP, coinciding with the establishment of the first sedentary hunter-gatherer settlements [22]. The earliest human-domesticated plants were grains [64], which also comprise a staple of wild mouse diets. However, commensalism was likely linked to an increased dietary reliance on grains and starch-rich foods, at the expense of seeds, fruits, insects, and other components of the wild mouse diet. This dietary shift potentially imposed strong selection to improve the efficiency of nutrient absorption from grains and starches. Indeed, we found clear evidence for recent positive selection at *Mgam*, a maltase-glucoamylase that plays a key role in the final stages of starch digestion. It is particularly noteworthy that signals of selection on this gene are limited to the mouse populations from Iran and Afghanistan, as these two populations coincide with some of the earliest human agricultural settlements [65] and overlap the presumed ancestral region of *M. musculus* [66, 67]. Strikingly, prior studies have also linked signals of positive selection at *Mgam* to the successful transition to agriculture in Andean human populations [42] and dietary shifts that accompanied the domestication of dogs [41]. We also identified a signal of selection near *Amy1* on chr3qF3 in the mouse population from France. Amylase is a presumed target of positive selection in human populations, with increased copy number linked to increased starch digestion capacity [68]. However, genetic adaptation in mice is likely rendered through short nucleotide variants, rather than copy number changes.

Fourth, many selective sweeps in wild house mice have occurred at genes that have been implicated in human diseases and disorders (Additional file 11: Table S7). Indeed, we show that targets of positive selection in wild mice are significantly encriched for disease-associated genes compared to null expectations. For example, multiple mouse populations harbor signals of selection at *Pkd2*, a gene that, when mutated in humans, can lead to polycystic kidney disease [48]. We also report selective sweeps spanning genes associated with autism spectrum disorder and speech-related impairment (e.g., *Grin2a, Cntnap2, Snrpn, Nrxn3, Herc2, Nalcn*, and *Cadps2*) [69, 70]. Understanding the mechanisms of adaptation at these genes in wild mouse populations could provide critical insights into the evolutionary basis of these diseases in humans.

In addition to these major themes, our analysis also presents a cautionary tale regarding the importance of integrating data on local genomic copy number with diversity metrics used in selection scans. Notably, several regions of significantly reduced diversity that emerged in our analysis proved to be false positives due to the presence of cryptic segregating structural variants. For example, a signal consistent with the positive selection at the *Skint* genes cluster on chr4:112.08 - 112.60 Mb in the Indian *M. m. castaneus* population is an artifact due to a high-frequency deletion spanning this region. While the joint analysis of copy number and genetic diversity metrics is not currently commonplace in genome-wide selection scans, our incidental findings argue that such practice should become routine.

Overall, our findings mirror conclusions for human populations, revealing that natural selection has shaped the geographic landscape of wild mouse variation in a manner that influences the distribution of likely disease-associated alleles. However, we note that our approach for identifying signals of positive selection is not designed to find signals of polygenic adaptation. In contrast to the hard selective sweep signatures reported here, wherein a single haplotype or variant is driven to high frequency within a population, signals of adapation on polygenic traits typically yield so-called “soft sweep” signatures, marked by milder increases in allele frequency of the high-fitness haplotype [71, 72]. Powerful approaches for detecting polygenic adaptation have been developed in recent years (e.g. [73]), and future efforts would be well spent by applying these methodologies to the wild mouse populations studied here.

### Conclusions

Successful adaptation to a commensal environment set the stage for subsequent human-aided dispersal of house mice across the globe, including the colonization of new environments in recent history. As a consequence of this demographic history and subsequent local adaptation, mice from different geographic regions are genetically and phenotypically differentiated, and notably at many loci associated with traits with immediate relevance to human health and disease. Overall, our analysis reveals that natural selection has played an important role in shaping global patterns of wild mouse diversity and spotlights key pathways and genes targeted by positive selection during recent house mouse evolutionary history. We anticipate that our catalog could help prioritize specific geographic areas for sampling wild mice to develop new natural mouse models of human disease or conduct genome-wide association studies in natural populations [7].

## Methods

### Whole-genome sequences

We analyzed a total of 154 previously published whole-genome sequences [6, 17, 18], including multiple populations from each of the three principle house mouse subspecies. In total, we surveyed four populations of *M. m. domesticus*, including 50 samples from the Eastern United States, 28 from France, six from mainland Germany, three from Heligoland, and seven from Iran. We analyzed 30 *M. m. castaneus* genomes from two populations (Taiwan, n = 20; India, n = 10), and 22 *M. m. musculus* genomes from three populations (Afghanistan, n = 6; Czech Republic, n = 8; Kazakhstan, n = 8). The sequence dataset also includes eight *M. spretus* genomes from Spain.

### Sequence alignment and variant calling

Fastq reads were mapped to the mm10 reference genome using the default parameters in BWA version 0.7.15 [74]. We followed the standard Genome Analysis Tool-kit (GATK; version 3.8.0) pipeline for subsequent pre-processing before variant calling [75, 76]. Variant calling was performed on each subspecies or species group using the “-ERC GVCF” mode in “HaplotypeCaller”. The “output” was then subjected to a series of hard filters using “-- filterExpression “QD < 2.0 ‖ FS > 60.0 ‖ MQ < 40.0 ‖ MQRankSum < -12.5 ‖ ReadPosRankSum < -8.0”. The resulting hard filtered variants were then used as training data for the “output” during the variant recalibration stage. Subsequent downstream analyses were restricted to biallelic autosomal variants.

### Variant annotation and statistics

We used SnpEff (version 4.3t) for both variant annotation and the determination of the total number of variants within each functional class per sample and per population [77]. The numbers of shared and unique variants between each subspecies and between species were calculated using the “vcf-stats” and “vcf-isec” commands within VCFtools (version 0.1.16) [78]. Variant sharing between taxonomic groups was visualized using the VennDiagram R package (version 1.6.20) [79].

### Assessing genetic relatedness

Closely related samples were identified using KING v.2.2.6 [80]. The full dataset includes 5 pairs of presumed first-degree relatives, 5 pairs of second-degree relatives, and 4 pairs putative third-degree relatives (Additional file 1: Table S1).

We used two approaches to assess levels of genetic relatedness among populations. We first thinned SNPs to one variant per 1 kb interval for all samples using VCFtools (version 0.1.16) [78] and then projected the data into two dimensions using a principal component analysis (Plink version 1.9) [81]. We also constructed a maximum likelihood phylogenetic tree from the 154 wild mouse genomes using PhyML (version 3.0) [82]. The best-fit nucleotide substitution model was determined using jModeltest (version 2.1.7) [83]. The resulting tree was visualized in MEGA (version 7) [84].

### Genome-wide copy number detection

Read depth was computed in 20 kb windows across each sequenced mouse genome using *mosdepth* [85]. Absolute read depth values were corrected for GC-content biases following established methods [86] and standardized by the genome-wide average read depth to convert to copy number (CN) estimates. Further, we retained only windows in diploid state but present in all the individual of each population using both the “awk” command and the “—intersect” option of the bedops version 2.4.39 [87]. We approximate the CN estimates on each window to its nearest whole number and exclude windows with CN less than or greater than two (or one for males on chrX) using the “intersect” option of bedtools version 2.29.2 [88].

### Identifying footprints of positive selection

As a beneficial allele increases in frequency under positive selection, it carries linked genetic variants with it, leaving behind a reduction in diversity at the targeted locus. To identify this signature of locally depressed diversity in the mouse genome, we computed three population genomic diversity statistics in 20 kb windows (10 kb sliding steps) across the genome: pool heterozygosity (*H*_p_) [30], nucleotide diversity (π) [31], and Tajima’s D [32]. Our analysis was restricted to variants on the autosomes and X chromosome, to the exclusion of Y-linked variants.

Windows with <50 SNPs were excluded, resulting in the elimination of ∼0.3% to ∼4% of all windows, depending on the population. Diversity statistics were normalized for each population to enable comparison across analyses. We used an empirical approach to identify putative regions of positive selection. Specifically, we focus on windows that reside at the extreme 0.1% tail of the observed *H*p distribution, but are supported by at least one of the π or Tajima’s D statistics. Although the computed statistics are not strictly independent of one another, they do encapsulate slightly different aspects of the patterning of genetic variation. Consequently, this requirement is likely to eliminate some false positives that present as outliers of the neutral distribution of diversity. Adjacent windows were then collapsed to form single candidate regions, similar to a previous study [89].

### Association with Mendelian traits and functional classification of putative sweep genes

We estimated the fraction of candidate sweep genes that overlap with genes in the OMIM database (https://www.omim.org/, retrieved October 22, 2020) and GWAS catalog (https://www.ebi.ac.uk/gwas/, accessed March 6, 2021). We then compared this fraction to the genome-wide null expectation using a simulation procedure. Briefly, we simulated a set of random genomic regions size-matched to the distribution of observed sweep windows and then computed the fraction of simulated regions harboring genes in the OMIM and GWAS database. We repeated this simulation procedure 1000 times to derive the expected frequency of human disease genes in sweep windows. We report all association with Mendelian traits in Additional file 11: Table S7.

For the functional classification, we retrieved genes within each candidate selective sweep region using Ensembl BioMart version 100 [90]. These gene lists, either at the population or subspecies level, were used for gene ontology and Kyoto Encyclopedia of Genes and Genomes enrichment analyses using the Database for Annotation, Visualization, and Integrated Discovery (DAVID version 6.8) [91]. Overrepresented gene clusters were identified by Fisher’s Exact tests (*P* < 0.05).

### Gene expression analyses

Publicly available transcriptome sequencing reads from 10 different tissues (gut, brain, heart, liver, lung, spleen, kidney, testis, thyroid, muscle) were obtained from wild-caught *M. m. domesticus* mice from Iran, France, and Germany [17]. Mapped reads were compiled into a count matrix using the featureCounts command in the Rsubread package. The resulting count matrix was then used to run a differential gene expression analysis across populations with the *edgeR* [92] and *DESeq2* [93] pipelines. The threshold for significance was set at p<0.01 in *edgeR* and adjP < 0.05 in DESeq2. Both methods produced largely overlapping sets of significantly differentially expressed genes across the populations.

Testis cell-type specific expression patterns for genes under positive selection on the X-chromosome were identified using the interactive mouse testis single cell RNAseq atlas [57].

## Declarations

### Ethics approval and consent to participate

Not applicable

### Availability of data and materials

All the genome sequences used in this study were obtained from three previously published studies [6, 17, 18].

### Consent for publication

Not applicable

### Contributions

R.A.L and B.L.D conceived of the project, performed analyses, and wrote the paper. U.P.A performed the differential gene expression analysis. All authors approved the final paper.

## Supplementary information

**Additional file 1: Table S1**. Relatedness among samples (.xlsx)

**Additional file 2: Figure S1**. Genome-wide distribution of positive selection signals in four populations of *M. m. domesticus*. The horizontal lines correspond to the 0.1% threshold from the tail end of the diversity statistics distribution. (.tiff)

**Additional file 3: Figure S2**. Genome-wide distribution of positive selection signals in two populations of *M. m. castaneus*. The horizontal lines correspond to the 0.1% threshold from the tail end of the diversity statistics distribution. (.tiff)

**Additional file 4: Figure S3**. Genome-wide distribution of positive selection signals in three populations of *M. m. musculus*. The horizontal lines correspond to the 0.1% threshold from the tail end of the diversity statistics distribution. (.tiff)

**Additional file 5: Table S2**. Candidate selective sweep regions, gene expression, and GO and KEGG analyses in the four populations of *M. m. domesticus*. (.xlsx)

**Additional file 6: Table S3**. Candidate selective sweep regions and GO and KEGG analyses in the three populations of *M. m. musculus*. (.xlsx)

**Additional file 7: Table S4**. Candidate selective sweep regions and GO and KEGG analyses in the two populations of *M. m. castaneus*. (.xlsx)

**Additional file 8: Table S5**. Candidate selective sweep regions and GO and KEGG analyses in *M. spretus*. (.xls)

**Additional file 9: Figure S4**. Cryptic structural variation at the *Skint* gene cluster (chr4:112.08 – 112.60 Mb) yields signals consistent with a selective sweep in the Indian *M. m. castaneus*. (**a**) ZHp for the two *M. m. castaneus* populations. The red horizontal line is the 0.1% threshold (ZHp = -3.9). (b,c) A high frequency deletion segregates at frequency of 0.9 in this population. Only one surveyed individual is diploid for this locus (**b**), with 20% heterozygous and 70% homozygous for the deletion haplotype (**c**). Panel (**d**) presents the organization of the *Skint* paralogs across this region. (.tiff)

**Additional file 10: Table S6**. Candidate selective sweep genes and testis cell-type expression profiles on the X chromosome across the 9 *M. musculus* populations and *M. spretus*. (.xlsx)

**Additional file 11: Table S7**. Candidate selective sweep genes and their association with Mendelian and complex traits. (.xls)

**Additional file 12: Figure S5:** RNA expression levels of *Amy1, Cry1, Epas1*, and *Mgam* in various tissues (A-J) collected from *M. m. domesticus* populations of Germany (GR), Iran (IR), and France (FR). RNA expression level is represented by log normalized counts of reads (y-axis) in the populations (x-axis). Genes highlighted in red have significant (Likelihood ratio test, adjP < 0.05) differential gene expression across populations in the particular tissue.

